# Myostatin is a negative regulator of adult neurogenesis in zebrafish

**DOI:** 10.1101/2021.08.18.456778

**Authors:** Vishnu Muraleedharan Saraswathy, Lili Zhou, Brooke Burris, Deepika Dogra, Sven Reischauer, Mayssa H. Mokalled

## Abstract

Intrinsic and extrinsic inhibition of axonal and neuronal regeneration obstruct spinal cord (SC) repair in mammals. In contrast, adult zebrafish achieve functional recovery after SC damage. While studies of innate SC regeneration have focused on axon regrowth as a primary repair mechanism, how local neurogenesis impacts functional recovery is unknown. We uncovered dynamic expression of *myostatin b* (*mstnb*) in a niche of dorsal ependymal progenitors after complete SC transection in zebrafish. Genetic loss-of-function in *mstnb* impaired functional recovery, although glial and axonal bridging across the lesion were unaffected. Using a series of transgenic reporter lines, we quantified the numbers of stem, progenitor, and neuronal cells in the absence of *mstnb*. We found neural stem cell proliferation was reduced, while newborn neurons were increased in *mstnb* null tissues, suggesting *mstnb* is a negative regulator of neurogenesis. Molecularly, neuron differentiation genes were upregulated, while the neural stem cell maintenance gene *fgf1b* was downregulated in *mstnb* mutants. Finally, we show that human FGF1 treatment rescued neuronal gene expression in *mstnb* mutants. These studies uncover unanticipated neurogenic functions for *mstnb* in adult zebrafish, and establish the importance of local neurogenesis for functional SC repair.

## INTRODUCTION

Traumatic spinal cord injury (SCI) causes irreversible neuronal and systemic deficits (Hachem et al., 2017; Silva et al., 2014; Singh et al., 2014). In mammals, axon regrowth and neurogenesis are impeded by intrinsic and extrinsic inhibitory mechanisms that obstruct spinal cord (SC) regeneration (Alizadeh et al., 2019; Oyinbo, 2011; Sofroniew, 2018; Tran et al., 2021). Although multiple cell types including astrocytes and oligodendrocyte progenitor cells proliferate after SCI, the mammalian SC is incapable of generating mature neurons *in vivo* (Horky et al., 2006; Horner et al., 2000; Yamamoto et al., 2001). In contrast with mammals, highly regenerative vertebrates including teleost fish spontaneously recover after SCI. Following complete transection of SC tissues, adult zebrafish extend glial and axonal bridges across the lesion and achieve functional recovery within 6 to 8 weeks of injury. In addition to axon regrowth from hindbrain neurons, zebrafish regenerate motor neurons and interneurons around the lesion site (Becker et al., 1998; Becker et al., 1997; Kuscha et al., 2012a; Mokalled et al., 2016; Reimer et al., 2008). Yet, the contribution of local neurogenesis to functional recovery and the mechanisms that coordinate the regeneration of different neuronal subtypes remain to be determined.

Ependymal radial glial cells (ERGs) line the brain ventricles and SC central canal. ERGs co-express astroglial (*gfap* and *blbp*) and progenitor (*sox2* and *hey*) cell markers, and comprise populations of neurogenic stem cells in adult zebrafish (Kroehne et al., 2011; März et al., 2010; Ogai et al., 2014; Reimer et al., 2009; Than-Trong et al., 2020; Than-Trong et al., 2018). In uninjured neural tissues, the majority of ERGs are non-dividing, quiescent cells that possess epithelial-like features (Barbosa et al., 2015; Chapouton et al., 2010; März et al., 2010). Following brain and SC damage, ERGs undergo widespread proliferation and are thought to act as a major source of regenerated neurons (Adolf et al., 2006; Barbosa et al., 2015; Grandel et al., 2006; Kroehne et al., 2011). Activated ERGs undergo 3 modes of cell division: 1) symmetric division into ERGs, 2) asymmetric division into ERG and neural progenitors, or 3) symmetric division to generate differentiated neurons (Barbosa et al., 2015; Rothenaigner et al., 2011). Distinct lineage-restricted ERG domains emerge during SC regeneration in zebrafish. ERGs within the progenitor motor neuron (pMN) domain express *olig2* and generate *isl1/2^+^* and *hb9^+^* motor neurons after SCI (Reimer et al., 2008). On the dorsal and ventral sides of the pMN, ERGs give rise to *vsx1^+^* V2 interneurons and serotonergic neurons, respectively (Barreiro-Iglesias et al., 2015; Kuscha et al., 2012b). We recently showed that ventral ERGs undergo epithelial-to-mesenchymal transition (EMT), and that EMT is required for glial bridging and functional regeneration (Klatt Shaw et al., 2021). Thus, despite their morphological similarities, ERGs elicit compartmentalized injury responses and proliferate into lineage-restricted progenitors during SC regeneration.

Myostatin (Mstn), also known as Growth differentiation factor 8 (Gdf8), is a Tgf-β superfamily member. Upon binding Activin type 2 and type 1 receptors, Mstn induces the phosphorylation and nuclear translocation of Smad 2/3 to regulate target gene expression (Massagué, 2012; Sartori et al., 2014; Sharma et al., 2015). Spontaneous and targeted Mstn loss-of-function mutations lead to double muscle phenotypes in zebrafish, mice, cattle and humans (Dogra et al., 2017; Kambadur et al., 1997; Schuelke et al., 2004; Whittemore et al., 2003). Mechanistically, Mstn controls lineage progression within myogenic stem cells (satellite cells) and progenitor cells (myoblasts). During muscle development, Mstn inhibits myoblast differentiation via negative regulation of the myogenic transcription factors MyoD and Myogenin. Mstn also suppresses satellite cell proliferation, differentiation, and muscle regeneration (Langley et al., 2002; McCroskery et al., 2003; McCroskery et al., 2005). However, it remains unclear whether recombinant MSTN proteins inhibit or stimulate myoblast proliferation (Rodgers et al., 2014; Taylor et al., 2001). Equally conflicting effects were reported for recombinant MSTN proteins on neuronal proliferation and neurite outgrowth *in vitro* (Kerrison et al., 2005; Wu et al., 2003), suggesting Mstn functions are dose- and context-dependent and that *in vivo* studies are required to decipher the role of Mstn in the nervous system.

Tgf-β signaling directs immune, fibrotic, or regenerative injury responses across tissues and species. As zebrafish SCs regenerate without fibrotic scarring, we postulated that Tgf-β signaling is pro-regenerative in adult zebrafish. By surveying the expression of Tgf-β ligands after SCI, we found *mstnb* is induced in dorsal ERGs of lesioned SC tissues. *mstnb* mutants showed normal baseline swim capacity but failed to achieve functional recovery following SCI, despite having normal axonal and glial bridging across the lesion. Mstnb inhibition using genetic loss-of-function and pharmacological approaches enhanced neurogenesis and diminished ERG proliferation. A series of transgenic reporter lines was used to quantify the numbers of neural stem cells (NSCs), intermediate neural progenitors (iNPs), and interneurons. These studies revealed NSC proliferation was reduced, while regenerating neurons were increased in *mstnb* mutants. RNA sequencing showed neuron differentiation genes were upregulated in *mstnb* mutants. Finally, we show that the neural stem cell maintenance gene *fgf1b* was downregulated in *mstnb* mutants, and that human FGF1 treatment rescued neuronal gene expression. These studies indicate that *mstnb* acts as an essential negative regulator of adult neurogenesis in zebrafish, and that injury-induced *mstnb* expression is required to maintain the potency and self-renewal of neurogenic ERGs during SC regeneration.

## RESULTS

### Tgf-β signaling is activated in dorsal ependymal progenitors after SCI

Injuries to the central and peripheral nervous systems induce Tgf-β signaling across vertebrates. In mammals, Tgf-β activation directs a range of regenerative and anti-regenerative cell responses including immune cell activation, neurite outgrowth, and scar formation (Li et al., 2017). In zebrafish larvae, the anti-inflammatory effects of Tgfb1a are required for SC regeneration (Keatinge et al., 2021). We postulated that Tgf-β signaling is pro-regenerative in adult zebrafish. To explore this hypothesis, we first surveyed Smad3 phosphorylation as a readout of Tgf-β activity after zebrafish SCI. By immunohistochemistry, phosphorylated Smad3 (pSmad3) was strongly induced in dorsal SC tissues at 1 week post-injury (wpi) (Fig. 1A). pSmad3 expression gradually diminished between 2 and 3 wpi, and was minimally expressed in uninjured SCs (Fig. 1A). At 1 wpi, pSmad3^+^ cells accounted for 7% of dorsal SC cells, and were reduced by 3 wpi relative to 1 wpi (Fig 1B, S1A). To determine the identity of Tgf-β responsive cells after SCI, we co-labelled pSmad3 with either the neuronal markers HuC and HuD (HuC/D) or the ependymal progenitor marker Sox2. At 1 wpi, we rarely observed vesicular pSmad3 expression in some HuC/D^+^ neurons. However, the majority of HuC/D^+^ neurons were pSmad3^-^ (Fig. S1B). pSmad3 was primarily expressed at high levels in Sox2^+^ ERGs (Fig. 1C). Quantification revealed 66 to 76% of pSmad3^+^ cells were Sox2^+^ across timepoints (Fig. 1D, S1C). By EdU incorporation, ∼10% of pSmad3^+^ cells were proliferative at 1 wpi in dorsal SCs, and pSmad3^+^ cell proliferation decreased to baseline levels by 3 wpi (Fig. S1D, E). These findings indicated Tgf-β signaling is activated in dorsal ERGs after SCI.

**Figure 1.**
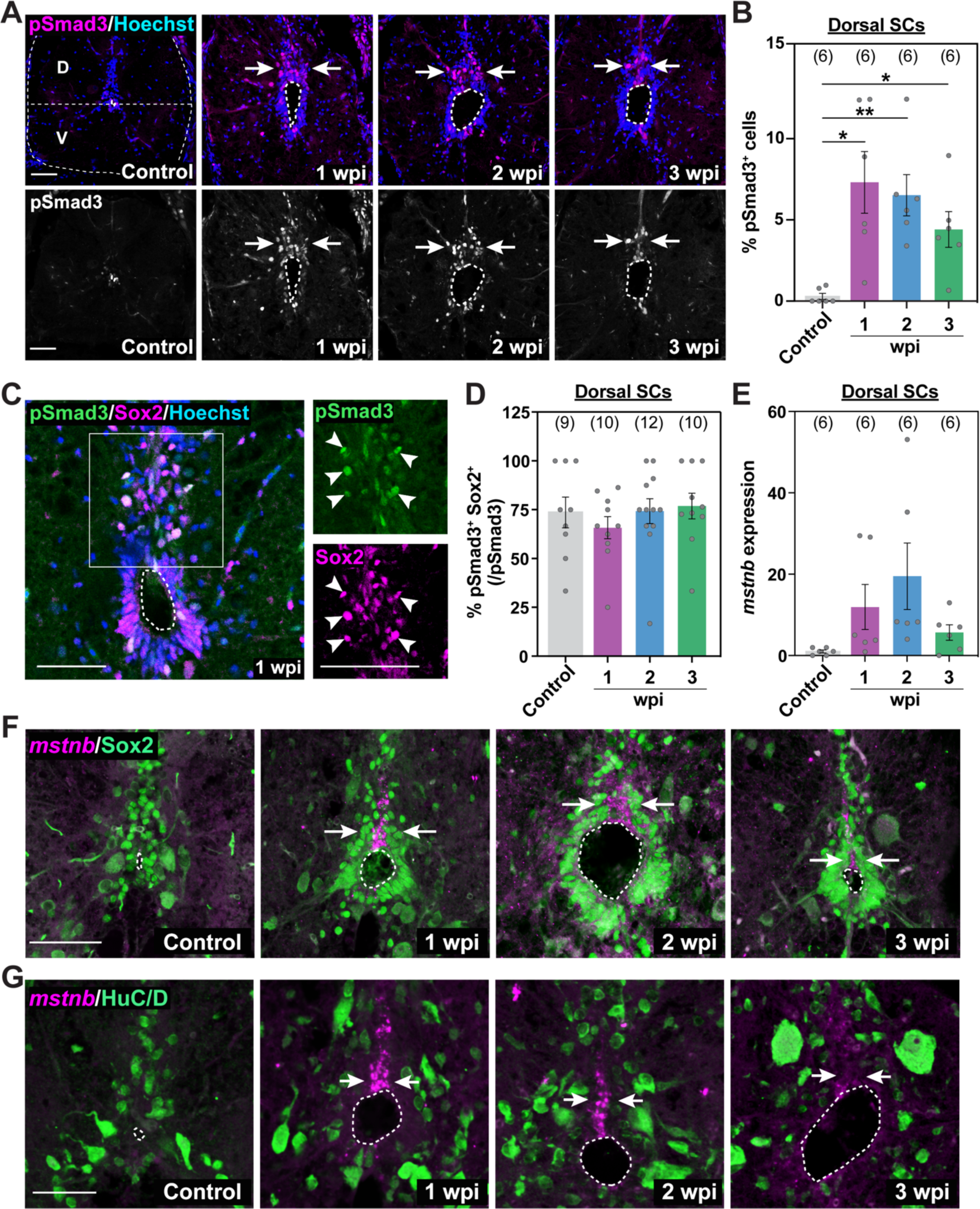
*mstnb* is induced in dorsal ependymal progenitors during SC regeneration. **(A)** Immunostaining for phosphorylated Smad3 (pSmad3) after SCI. Wild-type SC sections at 1, 2, and 3 wpi, and uninjured controls are shown. Cross sections 450μm from the lesion site are shown. Horizontal dotted line demarcates dorsal (D) and ventral (V) SC domains. Arrows point to pSmad3^+^ nuclei in the dorsal domain. **(B)** pSmad3 quantification in dorsal sections of wild-type SCs. Percent pSmad3^+^ cells was normalized to the number of nuclei in dorsal SCs. **(C)** pSmad3 and Sox2 immunostaining in wild-type SCs at 1 wpi. High-magnification views of dorsal SCs are shown. Arrowheads point to pSmad3^+^ Sox2^+^ ERGs. **(D)** pSmad3 quantification in dorsal ERGs. Percent pSmad3^+^ Sox2^+^ cells was normalized to the number of pSmad3^+^ cells. **(E)** Quantification of *mstnb* by *in situ* hybridization in dorsal SC tissues. **(F,G)** *mstnb* expression in wild-type SC sections after SCI. *mstnb* fluorescence *in situ* hybridization was followed by immunostaining for either Sox2 (F) or HuC/D (G) antibodies. Cross sections 450μm from the lesion site are shown at 1, 2, and 3 wpi, and for uninjured controls. Arrows point to domains of *mstnb* expression in dorsal SCs. Dotted ovals delineate central canal edges. For all quantification, SC sections 450 µm rostral to the lesion were analyzed and sample sizes are indicated in parentheses. *P<0.05; **P<0.01. Scale bars, 50 µm.

### *mstnb* expression is upregulated in dorsal ependymal progenitors after SCI

To explore mechanisms of Tgf-β activation during SC regeneration, we used a previously published RNA-seq dataset to survey the expression of Tgf-β ligands at 2 wpi (Mokalled et al., 2016). We found *mstnb*, *bmp2a/b*, and *bmp5* are upregulated, while *gdf3*, *gdf6a*, *gdf9*, *bmp6*, and *ndr1* are downregulated after injury relative to uninjured SC tissues (Fig. S1F). By fluorescence *in situ* hybridization, *mstnb* expression was induced in dorsal SC tissues between 1 and 2 wpi and decreased at 3 wpi (Fig. 1E-G). *mstnb* transcripts were not detectable in uninjured SCs (Fig. 1E-G). Co-labeling of *mstnb* transcripts with either ependymal Sox2 or neuronal HuC/D showed *mstnb* expression restricted to a subset of Sox2^+^ ERGs in dorsal SCs (Fig. 1F). *mstnb* transcripts were excluded from neuronal cell bodies (Fig. 1G). These studies revealed *mstnb* expression is induced in a subset of dorsal ERGs after SCI, and suggested *mstnb* expression correlates with Tgf-β activation in dorsal SC tissues during SC regeneration.

### *mstnb* is required for functional SC repair

To examine the role of *mstnb* during SC regeneration, we analyzed the extent of functional and cellular recovery in genetic zebrafish mutants (*mstnb^bns5^*) (Dogra et al., 2017) (Fig. 2A). *mstnb* mutants are adult viable, and elicit skeletal and cardiac muscle hyperplasia. Interestingly, the growth phenotypes associated with *mstnb* mutants are thought to be muscle specific (Dogra et al., 2017). To establish baseline motor function, we first assessed the swim capacities of wild-type, *mstnb* heterozygous (*mstnb^+/-^)* and homozygous (*mstnb^-/-^)* siblings in an enclosed swim tunnel under increasing water current velocities (Klatt Shaw and Mokalled, 2021; Klatt Shaw et al., 2021; Mokalled et al., 2016) (Fig. 2B). In this swim endurance assay, wild-type animals swam for 41 min before reaching exhaustion. *mstnb^+/-^* and *mstnb^-/-^* fish showed comparable swim functions, averaging 43 and 39 min of swim time, respectively. These results indicated *mstnb* mutants show normal swim capacity and suggested the muscle phenotype of *mstnb* mutants does not impact swim endurance. Next, we performed SC transections on *mstnb^-/-^* and control siblings and evaluated their functional regeneration between 2 and 6 wpi. *mstnb^+/-^* fish displayed normal swim capacity at 2 and 4 wpi, but their functional recovery was slightly compromised at 6 wpi (Fig. 2C). Relative to wild-type controls, functional recovery was 50% reduced in *mstnb^-/-^* fish at 2, 4, and 6 wpi (Fig. 2C).

**Figure 2.**
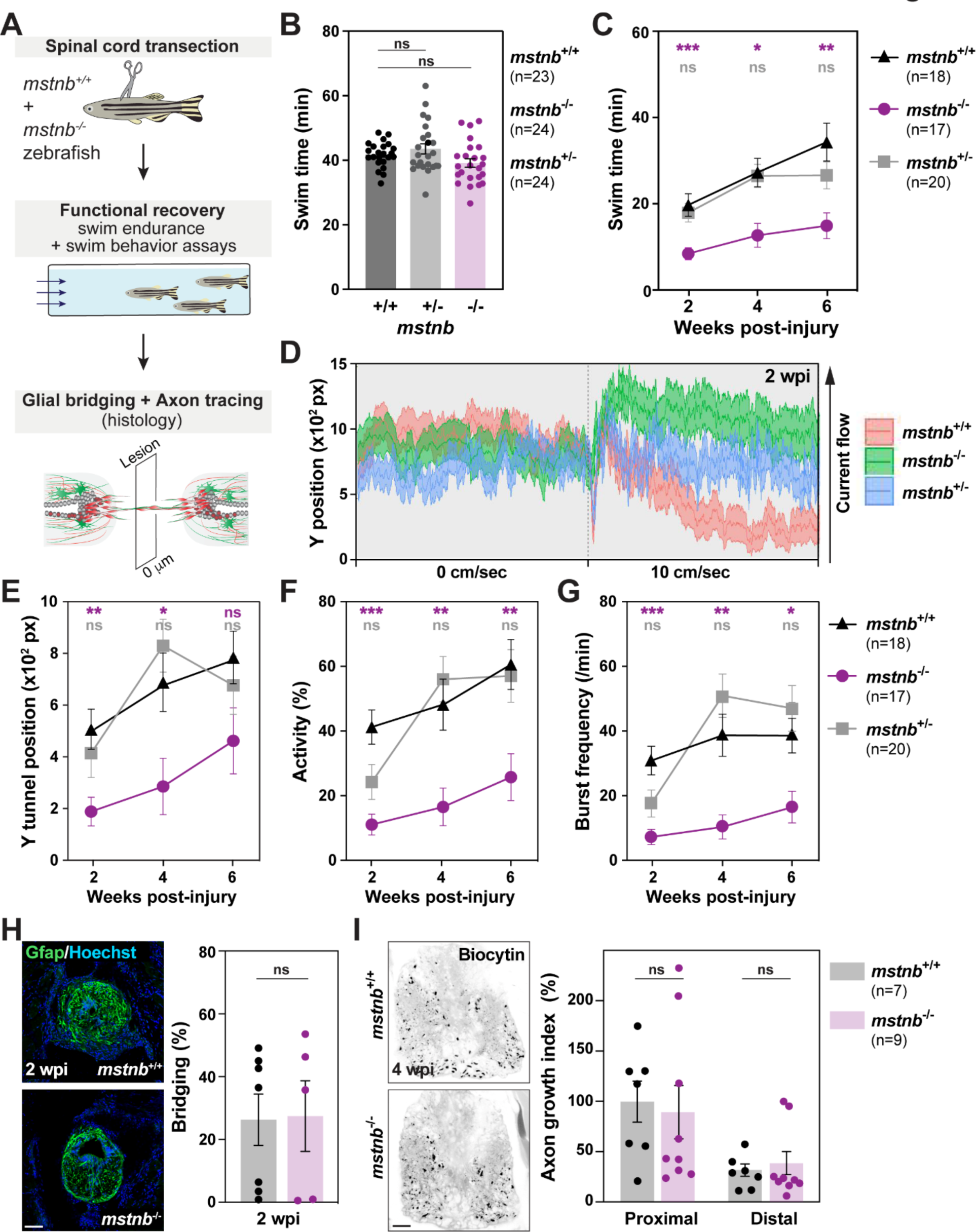
*mstnb* is required for functional SC regeneration. **(A)** Experimental pipeline to examine regeneration phenotypes after SCI. *mstnb^-/-^* fish and wild-type siblings were subjected to complete SC transection. Functional recovery was assessed between 2 and 6 wpi. Histology was used to assess glial and axonal bridging at 2 and 4 wpi, respectively. **(B)** Endurance swim assays determined baseline motor function for *mstnb^-/+^*, *mstnb^-/-^*, and wild-type fish. Dots represent individual animals from three independent clutches. **(C)** Endurance swim assays for *mstnb^-/+^*, *mstnb^-/-^*, and wild-type fish at 2, 4, and 6 wpi. Dots represent individual animals from two independent experiments. Statistical analyses of swim times are shown for *mstnb^-/+^* (grey) and *mstnb^-/-^* (magenta) relative to wild types. Recovery of *mstnb^-/-^* animals was not significant between 2 and 6 wpi. **(D)** Tracking swim performance at minimal water current velocity for *mstnb^-/+^* (blue), *mstnb^-/-^* (green), and wild-type siblings (red). Average Y position is shown for each cohort at 2 wpi. Animals were tracked in the absence of current (0 cm/sec) for 5 min, and for a 10cm/sec current velocity for 5 min. The arrow shows the direction of the water current. **(E-G)** Average Y position in the tunnel (E), percent activity (F), and burst frequency (G) were quantified at 10 cm/sec water current velocity. *mstnb^-/+^*, *mstnb^-/-^*, and wild-type fish are shown at 2, 4, and 6 wpi. Statistical analyses of swim times are shown for *mstnb^-/+^* (grey) and *mstnb^-/-^* (magenta) relative to wild types. Two independent experiments are shown. **(H)** Glial bridging in *mstnb^-/-^* (magenta) and wild-type siblings (grey) at 2 wpi. Representative immunohistochemistry shows the Gfap^+^ bridge at the lesion site. Percent bridging represents the cross-sectional area of the glial bridge at the lesion site relative to the the intact SC. Percent bridging was quantified for 7-9 animals per group. **(I)** Anterograde axon tracing in in *mstnb^-/-^* (magenta) and wild-type zebrafish (grey) at 4 wpi. Biocytin axon tracer was applied rostrally and analyzed at 100 μm (proximal) and 500 μm (distal) caudal to the lesion. Representative traces of biocytin are shown for each genotype animals at the proximal level. Quantification represents 7-9 animals per group. Axon growth was normalized to Biocytin labeling in wild-types at the proximal level. *P<0.05; **P<0.01; ***p<0.001; ns, not significant. Scale bars, 50 µm.

To further rule out the contribution of skeletal muscle overgrowth to the functional regeneration output of *mstnb* mutants, we tracked the swim behavior of *mstnb* mutants in the absence of water current or under a constant, low current velocity of 10 cm/sec (Fig. 2D). We reasoned that, unlike the endurance test that required fish to swim against increasing current velocities, swim behavior under minimal current velocity is less likely to be dependent on muscle function. Fish position in the swim tunnel (Y position), percent activity, and burst frequency were quantified to assess overall swim competence. In this assay, *mstnb^-/-^* animals stalled in the back quadrant of the swim tunnel (Fig. 2E), were 65% less active than their wild-type siblings (Fig. 2F), and displayed less bursts under low current velocity (Fig. 2G). Consistent with a partial regeneration phenotype in heterozygous fish, *mstnb^+/-^* fish were 40% less active and their burst frequency was reduced by 35% relative to wild types at 2 wpi. Swim parameters were comparable between *mstnb^+/-^* and wild-type siblings at 4 and 6 wpi, and were not statistically significant at 2 wpi (Fig. 2E-G).

To date, cellular growth across the lesion site has served as a primary readout of cellular regeneration in zebrafish (Goldshmit et al., 2012; Mokalled et al., 2016; Reimer et al., 2013). Glial bridging and axon tracing assays were performed to evaluate the extents of glial and axonal regeneration across the lesion site (Fig. 2A). By Gfap immunostaining, glial bridging was unaffected in *mstnb^-/-^* animals compared to wild-type siblings at 2 wpi (Fig. 2H). At 4 wpi, anterograde axon tracing using Biocytin showed comparable axon regrowth in proximal and distal SC sections between *mstnb^-/-^* and control animals (Fig. 2I). Together, these studies indicated *mstnb* is required for functional SC repair but is dispensable for glial bridging and axonal regrowth across the lesion. These findings prompted us to investigate mechanisms of SC regeneration that are independent of glial bridging and axon growth.

### *mstnb* is a negative regulator of adult neurogenesis after SCI

In addition to glial bridging and axon regrowth, zebrafish regenerate lost motor neurons and interneurons around the lesion site (Barreiro-Iglesias et al., 2015; Kuscha et al., 2012b; Reimer et al., 2008). Dorsal ependymal progenitors are thought to give rise to regenerating interneurons in dorsal SC tissues after injury. Since *mstnb* is expressed in dorsal ERGs after SCI and *mstnb* mutants did not show glial bridging or axon growth defects, we postulated that *mstnb* plays a role in adult neurogenesis in zebrafish and that local neurogenesis around the lesion is required for functional SC repair. To test this hypothesis, we first examined the proliferation rates of Sox2^+^ ERGs and of regenerating HuC/D^+^ neurons in *mstnb* mutants. Uninjured and injured *mstnb^-/-^* and wild-type siblings were subjected to SCI and to a single EdU pulse 24 hours prior to SC collection and histological analysis (Fig. 3A and S2A). Cell counts revealed a significant increase in HuC/D^+^ EdU^+^ neurons in dorsal SCs with 1 wpi SCs showing the most pronounced differences (Fig. 3B, C and S2B). At 1 wpi, 7.7% of HuC/D^+^ neurons were EdU^+^ in wild-type SCs (Fig. S2B), and accounted for 0.9% of dorsal SC cells (Fig. 3C). Conversely, 15.2% of HuC/D^+^ neurons were EdU^+^ in *mstnb^-/-^* SCs (Fig. S2B), accounting for 1.5% of dorsal SC cells (Fig. 3C). The rates of neurogenesis where attenuated in wild-type SCs at 2 and 3 wpi relative to 1 wpi. However, *mstnb* mutants showed increased neurogenesis at 3 wpi (Fig. 3C and S2B). These differences were blunted in cell counts from total SCs (Fig. S2C), suggesting neuronal differentiation is specifically increased in dorsal SC tissues of *mstnb* mutants. On the other hand, we observed a minor, non-significant decrease in the number of Sox2^+^ EdU^+^ ERGs in *mstnb^-/-^* SCs at 1 and 3 wpi (Fig. 3D, E and S2D, E). These findings indicated the rates of neurogenesis are increased in *mstnb* mutants.

**Figure 3.**
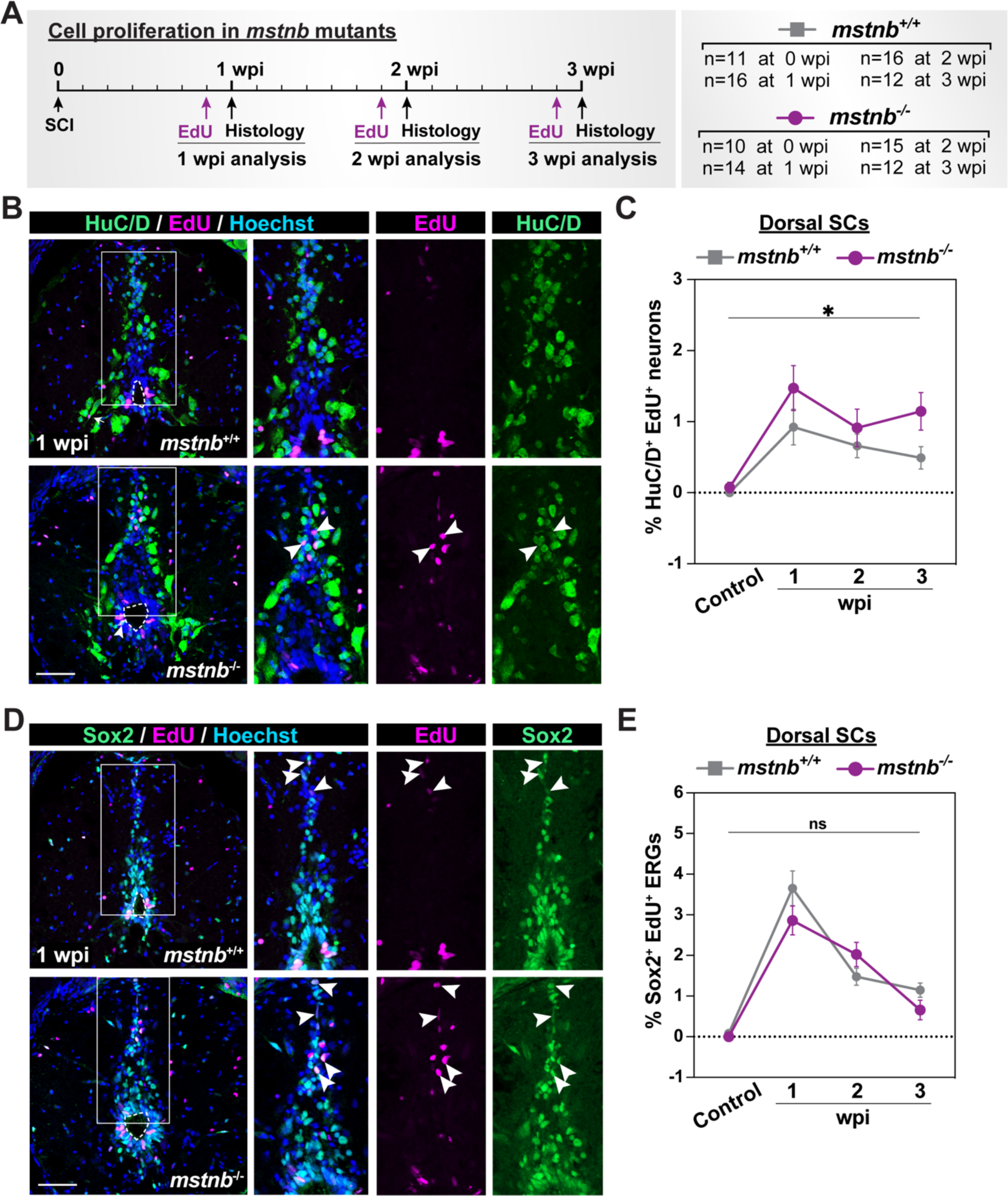
Cell proliferation in *mstnb* mutant zebrafish. **(A)** Experimental timeline to assess the rates of cell proliferation. *mstnb^-/-^* and wild-type siblings were subjected to SC transections. A single EdU injection was performed at either 6, 13, or 20 days post-injury. SC tissues were harvested for analysis at 1, 2, 3 wpi and at 24 hours after EdU injection. Animal numbers are indicated for each genotypes and two independent replicates are shown. **(B)** Immunohistochemistry for EdU and HuC/D in SC sections of *mstnb^+/+^* and *mstnb^-/-^* at 1 wpi. The region inside the rectangular box is shown in higher magnification. Arrowheads indicate HuC/D^+^ EdU^+^ neurons. **(C)** Regenerated HuC/D^+^ EdU^+^ neurons were quantified in dorsal SC sections at 1, 2, 3 wpi and uninjured controls. Percent HuC/D^+^ EdU^+^ neurons was normalized to the total number of nuclei for each section. **(D)** Immunohistochemistry for EdU and Sox2 in SC sections of *mstnb^+/+^* and *mstnb^-/-^* at 1 wpi. The region inside the rectangular box is shown in higher magnification. Arrowheads indicate Sox2^+^ EdU^+^ ERGs. **(E)** Sox2^+^ EdU^+^ ERGs were quantified in dorsal SC sections at 1, 2, 3 wpi and uninjured controls. Percent Sox2^+^ EdU^+^ ERGs was normalized to the total number of nuclei for each section. For all quantifications, cross SC sections 450 μm rostral to the lesion site were quantified.*P<0.05; ns, not significant. Scale bars, 50 µm. Dotted ovals delineate central canal edges.

To evaluate how snapshots of increased neurogenesis at 1 wpi could impact the numbers of regenerating neurons in *mstnb* mutants, we performed SC transections on *mstnb^-/-^* and wild-type fish followed by daily EdU injections for 1 or 2 wpi (Fig. 4A and S3A). In this assay, daily EdU labeling allowed us to estimate the total numbers of regenerating neurons (HuC/D^+^ EdU^+^ neurons), and the extent of self-renewal in ependymal progenitors (Sox2^+^ EdU^+^ ERGs). Considering the dynamic rates of neurogenesis along the rostro-caudal axis, we quantified cell numbers at 150, 450, and 750 μm rostral to the lesion. At 1 wpi and in wild-type sections proximal to the lesion site (-150 μm), 10.9% of dorsal SC cells were HuC/D^+^ neurons (Fig. 3C), and 2.7% of dorsal SC cells were regenerating neurons (HuC/D^+^ EdU^+^) (Fig. 3D). On the other hand, 19% of dorsal SC cells were HuC/D^+^ neurons (Fig. 3C), and 4.3% of dorsal SC cells were regenerating neurons (HuC/D^+^ EdU^+^) in *mstnb^-/-^* SCs (Fig. 3D). Quantifications from total SC tissues confirmed that the increase in regenerating neurons in *mstnb* mutants is specific to dorsal SCs and blunted in total SCs (Fig. S3B,C). At 2 wpi, the numbers of HuC/D^+^ EdU^+^ neurons continued to be elevated in *mstnb^-/-^* SCs (Fig. 4D). Intriguingly, the overall increase in the total numbers of neurons was less pronounced at this time point (Fig. 4C), suggesting *mstnb* mutants elicit an early wave of increased neurogenesis at 1 wpi, but that compensatory mechanisms may be activated at later time points to counteract the loss of *mstnb*. Finally, despite increased neurogenesis rates in *mstnb^-/-^* fish, the numbers of Sox2^+^ and Sox2^+^ EdU^+^ ERGs were comparable across genotypes (Fig. 4F,G), suggesting ERG self-renewal was maintained at normal levels in *mstnb* mutants. These studies revealed a 2-fold increase in regenerating neurons in *mstnb* mutants, and are consistent with *mstnb* acting as a negative regulator of adult neurogenesis in zebrafish.

**Figure 4.**
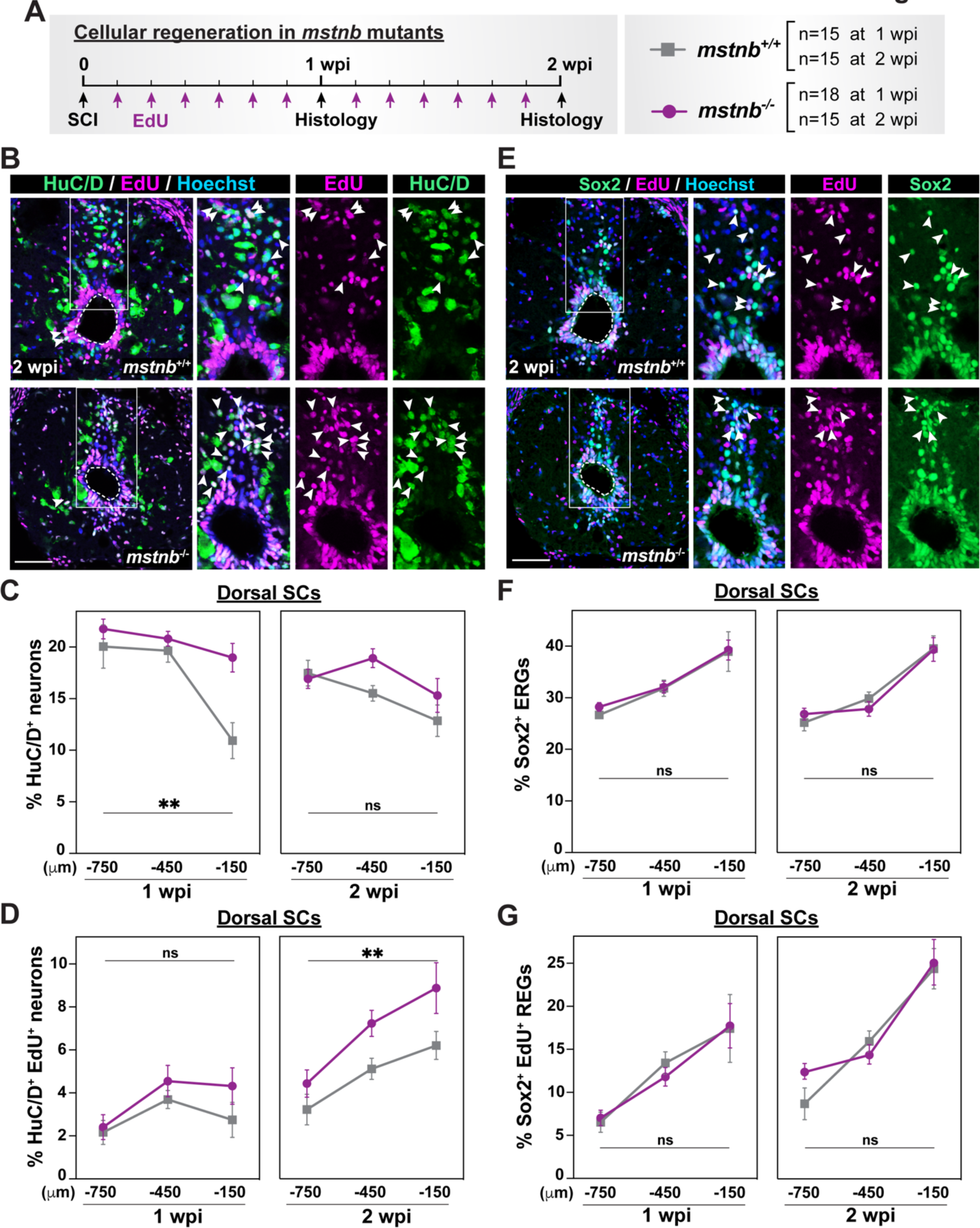
Regenerative neurogenesis in *mstnb* mutant zebrafish. **(A)** Experimental timeline to assess the rates of neurogenesis and ERG self-renewal. *mstnb^-/-^* and wild-type siblings were subjected to SC transections and daily EdU injections. SC tissues were harvested for analysis at 1 or 2 wpi. Animal numbers are indicated for each genotypes and two independent replicates are shown. **(B)** Immunohistochemistry for EdU and HuC/D in SC sections of *mstnb^+/+^* and *mstnb^-/-^* at 2 wpi. The region inside the rectangular box is shown in higher magnification. Arrowheads indicate HuC/D^+^ EdU^+^ neurons. **(C)** HuC/D^+^ neurons were quantified in dorsal SC sections at 1 and 2 wpi. Percent HuC/D^+^ neurons was normalized to the total number of nuclei for each section. **(D)** Regenerated HuC/D^+^ EdU^+^ neurons were quantified in dorsal SC sections at 1 and 2 wpi. Percent HuC/D^+^ EdU^+^ neurons was normalized to the total number of nuclei for each section. **(E)** Immunohistochemistry for EdU and Sox2 in SC sections of *mstnb^+/+^* and *mstnb^-/-^* at 2 wpi. The region inside the rectangular box is shown in higher magnification. Arrowheads indicate Sox2^+^ EdU^+^ ERGs. **(F)** Sox2^+^ ERGs were quantified in dorsal SC sections at 1 and 2 wpi. Percent Sox2^+^ ERGs was normalized to the total number of nuclei for each section. **(G)** Sox2^+^ EdU^+^ ERGs were quantified in dorsal SC sections at 1 and 2 w. Percent Sox2^+^ EdU^+^ ERGs was normalized to the total number of nuclei for each section. For all quantifications, cross SC sections at 150, 450, and 750 μm rostral to the lesion site were quantified. **P<0.01; ns, not significant. Scale bars, 50 µm. Dotted lines delineate central canal edges.

Human MSTN proteins are translated as inactive full-length precursors that undergo proteolytic processing into mature MSTN peptide and MSTN proform (pro-MSTN) peptide. pro- MSTN exhibits high binding affinity for Myostatin and inhibits its function (Zhu et al., 2000). To examine whether the global effects of *mstnb* mutants could be reproduced by local Mstnb inhibition, we injured wild-type animals and performed daily injections of human recombinant MSTN Proform (pro-MSTN) peptide adjacent to the lesion site (Fig. S4A). We then assessed the numbers of HuC/D^+^ neurons and Sox2^+^ ERGs at 1 wpi, corresponding to 6 days after initial treatment. We found HuC/D^+^ neurons were increased by 12% upon pro-MSTN treatment, though these differences were not significant (Fig. S4B). On the other hand, the numbers of Sox2^+^ progenitors were decreased by 20% in pro-MSTN injected fish relative to vehicle controls (Fig. S4C). Consistent with genetic loss-of-function of *mstnb*, pharmacological Mstn inhibition at the lesion site disrupted the relative ratios of HuC/D^+^ neurons and Sox2^+^ ERGs towards increased neurogenesis.

### *mstnb* mutants exhibit increased neuronal differentiation after SCI

Our proliferation assays showed increased neurogenesis and suggested the rate of ERG self- renewal was slightly decreased in the absence of *mstnb*. To dissect the cellular basis for this phenotype, we evaluated the numbers of neural stem cells (NSCs) and intermediate neural progenitors (iNPs) in *mstnb* mutants at baseline, 1 and 2 wpi (Fig. 5A). We first combined a *nestin* reporter transgene (*nes:*GFP) with *mstnb^-/-^* background to quantify NSCs in *nes:*GFP;*mstnb^-/-^* fish (Lam et al., 2009) (Fig. 5B-D and S5A-C). *nes*:GFP^+^ NSCs were rarely identified in uninjured SC sections from either mutant or wild-type animals, but were readily detectable after SCI (Fig. 5C and S5A). The proportions of *nes:*GFP^+^ cells in dorsal or total SC tissues were comparable between *mstnb^-/-^* and wild-type siblings at either 1 or 2 wpi (Fig. 5C and S5A). At these time points, NSC proliferation showed decreasing trends in *mstnb* mutants (Fig. 5D), and was statistically significant when the numbers of *nes:*GFP^+^ PCNA^+^ cells were normalized to *nes:*GFP^+^ NSCs (Fig. S5C). NSC proliferation was less pronounced in quantifications from total SC sections (Fig. S5B). These findings revealed NSC proliferation is reduced in *mstnb* mutants at 1 wpi.

**Figure 5.**
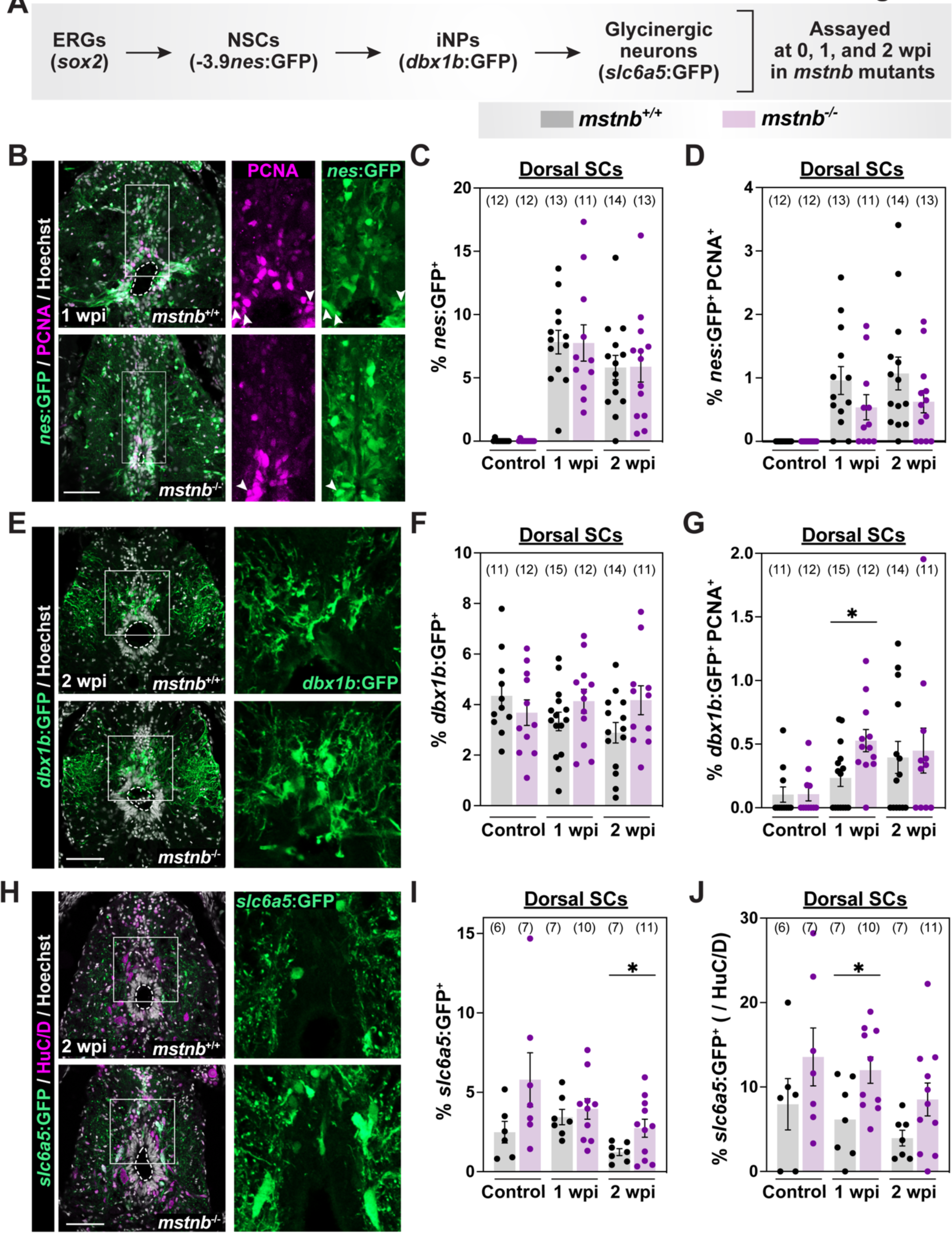
Assessment of neuronal progenitors and neurons in *mstnb* mutant zebrafish. **(A)** Experimental timeline to elucidate the dynamics of neurogenesis. *mstnb^-/-^* animals were crossed into either *-3.9nestin*:GFP, *dbx1b*:GFP, or *slc6a5*:GFP to evaluate the numbers of neural stem cells (NSCs), intermediate neural progenitors (iNPs), and glycinergic neurons, respectively. *mstnb^-/-^* and wild-type siblings were subjected to SC transections and collected at 1 or 2 wpi for analysis. Uninjured controls were used. Animal numbers are indicated for each genotypes and two independent replicates are shown. **(B)** GFP and PCNA staining in *-3.9nestin*:GFP*;mstnb^-/-^* SC sections at 1 wpi. *-3.9nestin*:GFP*;mstnb^+/+^* siblings are used as controls. The region inside the rectangular box is shown in higher magnification. **(C)** *nes^+^* NSCs were quantified in dorsal SC sections. Percent *nes^+^* NSCs was normalized to the total number of nuclei for each section. **(D)** *nes^+^* PCNA^+^ NSCs were quantified in dorsal SC sections. Percent *nes^+^* PCNA^+^ NSCs was normalized to the total number of nuclei for each section. **(E)** GFP staining in *dbx1b*:GFP*;mstnb^-/-^* SC sections at 2 wpi. *dbx1b*:GFP*;mstnb^+/+^* siblings are used as controls. The region inside the rectangular box is shown in higher magnification. **(F)** *dbx1b^+^* iNPs were quantified in dorsal SC sections. Percent *dbx1b^+^* iNPs was normalized to the total number of nuclei for each section. **(G)** *dbx1b^+^* PCNA^+^ NSCs were quantified in dorsal SC sections. Percent *dbx1b^+^* PCNA^+^ iNPs was normalized to the total number of nuclei for each section. **(H)** GFP staining in *slc6a5*:GFP:GFP*;mstnb^-/-^* SC sections at 2 wpi. *slc6a5*:GFP:GFP*;mstnb^+/+^* siblings are used as controls. The region inside the rectangular box is shown in higher magnification. **(I)** *slc6a5*:GFP*^+^* glycinergic neurons were quantified in dorsal SC sections. Percent *slc6a5*:GFP*^+^* neurons was normalized to the total number of nuclei for each section. **(J)** Percent *slc6a5*:GFP*^+^* neurons was normalized to the numbers of HuC/D^+^ neurons for each section. Dotted ovals delineate central canal edges. For all quantifications, cross SC sections 450 μm rostral to the lesion site were quantified. *P<0.05; ns, not significant. Scale bars, 50 µm.

The number of *nes*:GFP^+^ NSCs were unaltered in *mstnb* mutants despite their decreased proliferation rate. We postulated that *mstnb* loss-of-function may bias NSC fate towards neuronal differentiation, and that compensatory mechanisms upstream of NSC activation maintain their total numbers across genotypes. To test this hypothesis, we examined the number of iNPs using *dbx1b:*GFP transgene bred into a *mstnb^-/-^* background (Pierani et al., 2001; Satou et al., 2012) (Fig. 5E-G and S5D,E). The proportion of *dbx1b:*GFP^+^ iNPs averaged between 2.9 and 4.3 % of dorsal SC cells across time points, and showed an elevated trend in *mstnb^-/-^* relative to wild-type siblings at 2 wpi (Fig. 5F and S5D). These phenotypes were more pronounced in quantifications of *dbx1b:*GFP^+^ PCNA^+^ iNPs. Relative to wild-type siblings, *mstnb^-/-^* iNP proliferation was minimal prior to injury, increased by 2.3-fold at 1 wpi, and was normalized to wild-type levels by 2 wpi (Fig. 5G and S5E). These findings are consistent with accelerated neurogenesis in *mstnb* mutants.

To examine the differentiation and relative distribution of iNP-derived neurons after SCI, we labeled glycinergic neurons in *msntb* mutants using *slc6a5*:GFP reporter line (Fig. 5H-J and S5F) (McLean et al., 2007). At 1 wpi, the proportions of glycinergic neurons comprised 3.9% and 3.4% of dorsal SC cells in *mstnb^-/-^* and wild-type fish, respectively (Fig. 5I and S5F). By 2 wpi, glycinergic neurons accounted for 1.2% of dorsal SC cells in wild-type controls, but increased to 2.7% of dorsal cells in *mstnb^-/-^* SCs (Fig. 5I and S5F). By quantifying the proportions of glycinergic neurons within HuC/D^+^ neurons, glycinergic neurons accounted for 6.1% and 3.7% of neurons in wild-type SC tissues at 1 and 2 wpi, respectively (Fig. 5J). These proportions were 2-fold elevated in *mstnb* mutants, accounting for 12% of neurons at 1 wpi and 8.5% of neurons at 2 wpi (Fig. 5J). Together, these results indicated *mstnb* mutants exhibit an expansion in glycinergic neurons, which are overrepresented relative to other neuronal cell populations.

### Neuronal genes are upregulated in *mstnb* mutants

To determine the molecular mechanisms by which *mstnb* regulates the rates of neurogenesis, we deep sequenced SC tissues from *mstnb^-/-^* and wild-type siblings at 1 wpi, as well as uninjured controls (Fig. 6A-C). Principle component analysis confirmed clustering of biological replicates, and highlighted four distinct molecular signatures that are both injury- and genotype-induced (Fig. 6A). At 1 wpi, 61 genes were downregulated and 359 genes were upregulated in *mstnb^-/-^* SCs, suggesting *mstnb* may be a negative regulator of gene expression after SCI (Fig. 6C). Genes upregulated in *mstnb^-/-^* SCs comprised several neuronal or neuron differentiation genes, including *birc5b*, *eloal*, *elavl2*, *htr2aa*, and *pou5f3*, which were either unchanged or downregulated in uninjured *mstnb^-/-^* SCs (Fig. 6B-D). These findings indicated neuronal gene expression changes in *mstnb* mutants are injury-dependent.

**Figure 6.**
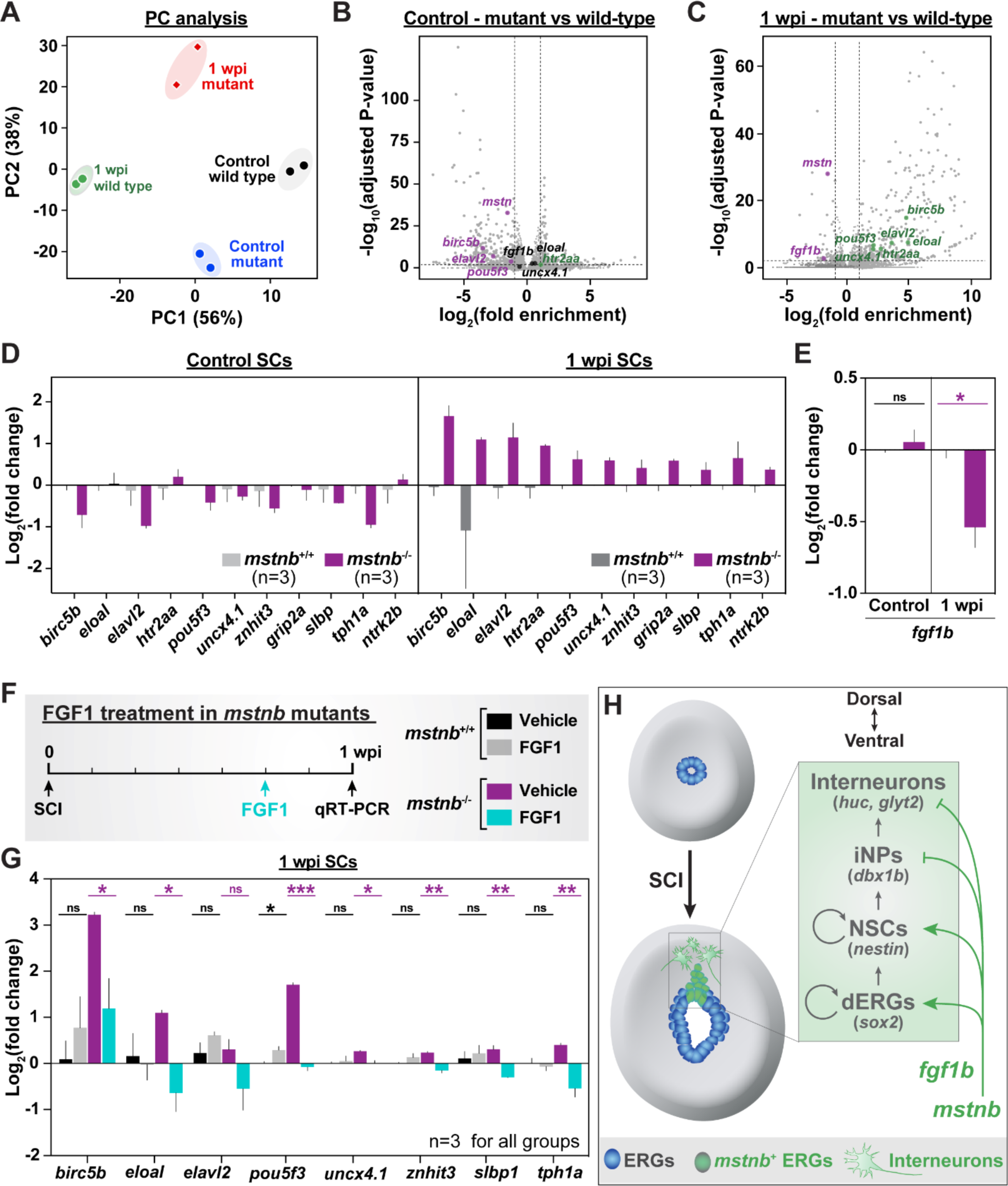
*mstnb* regulates neuronal gene expression via *fgf1b*. **(A)** *mstnb^-/-^* and wild-type siblings were subjected to complete SC transections and collected at 1 wpi for RNA sequencing. Control SC tissues were collected from uninjured fish. Sequencing was performed in independent duplicates. Principle component analysis shows clustering of biological replicates. Principle components 1 and 2 (PC1 and PC 2) show 56% and 38% variance, respectively. **(B,C)** Volcano plot representation of genes that are significantly upregulated or downregulated or depleted in *mstnb^-/-^* SCs relative to wild-type controls. Upregulated genes included genes with log2 (fold enrichment) > 1 and adjusted P-value <0.01. Downregulated genes included genes with log2 (fold enrichment) < -1 and adjusted P-value <0.01 are considered downregulated. Select upregulated (green) and downregulated (magenta) neuronal genes are indicated. Unchanged genes are labelled in black. **(D)** qRT-PCR for neuronal genes was performed on *mstnb^-/-^* and wild-type SCs at 1 wpi. Uninjured *mstnb^-/-^* and wild-type controls were used. For each time point, log2 (fold change) was normalized to *eif1α* and to gene expression levels in *mstnb^+/+^* controls. **(E)** *fgf1b* qRT-PCR was performed on uninjured and injured *mstnb^-/-^* and wild-type animals. For each time point, *fgf1b* expression was normalized to *eif1α* as a loading control and to *fgf1b* levels in *mstnb^+/+^* controls. **(F)** *mstnb^-/-^* and wild-type siblings were subjected to SC transections and treated with gelfoam-soaked human recombinant FGF1 at 5 dpi. *mstnb^-/-^* and wild-type controls were treated with vehicle-soaked gelfoam. SCs were collected for gene expression analysis at 1 wpi, which corresponds to 2 days post-treatment. Animal numbers are indicated for each genotype. **(G)** Neuronal gene expression was analyzed by qRT-PCR from FGF1- and vehicle-treated SC tissues. For each gene, log2 (fold change) was normalized to *eif1α* as a loading control and to gene expression levels in uninjured *mstnb^+/+^* SCs. **(H)** Schematic model shows *mstnb* is a negative regulator of neuronal differentiation in dorsal SC tissues after spinal cord injury. ERGs (blue) undergo extensive proliferation after SCI. A *mstnb* expressing niche (green) emerges in the dorsal ependyma.

Fibroblast growth factor (Fgf) maintains the proliferation and self-renewal capacities of neural stem cells in mammals (Hsu et al., 2009). By RNA-seq and qRT-PCR, *fgf1b* was downregulated in *mstnb^-/-^* SCs at 1wpi, but was unchanged in uninjured SC tissues (Fig. 6E). The dysregulation of *fgf1b* in *mstnb* mutants suggested *mstnb*-mediated *fgf1b* expression inhibits neurogenesis by promoting progenitor cell proliferation and self-renewal. To test this hypothesis, we examined whether the neuronal gene expression changes observed in *mstnb* mutants could be rescued by localized delivery of human recombinant FGF1 into SC lesions. We injured *mstnb^-/-^* and wild-type siblings and applied FGF1 proteins adjacent to the lesion site at 5 dpi using a gelfoam sponge. Gene expression changes were assessed by qRT-PCR at 1 wpi, corresponding to 2 days after treatment (Fig. 6F). Consistent with increased neurogenesis in *mstnb* mutants, *birc5b*, *eloal*, and *pou5f3* transcript levels were increased in vehicle-treated *mstnb^-/-^* relative to vehicle-treated wild- types (Fig. 6G). Application of exogenous FGF1 proteins at the lesion rescued the upregulation of neuronal genes in *mstnb^-/-^* SCs (Fig. 6G). These findings indicated Mstn-mediated Fgf signaling is a negative regulator of adult neurogenesis after zebrafish SCI.

## DISCUSSION

This study shows *mstnb* expression is induced in a subset of dorsal ERGs after SCI. Our results are consistent with a model in which *mstnb* regulates the rates of self-renewal and neuronal differentiation after SCI, and suggest *mstnb-*dependent Fgf signaling promotes self-renewal at the expense of neurogenesis (Fig. 6H).

Successful SC regeneration requires faithful recovery of the excitatory and inhibitory (E/I) balance in regenerating neural circuits. SCI alters the amount, strength, and relative locations of E/I inputs by disrupting descending hindbrain connections and promoting waves of axonal degeneration, neuronal death, and demyelination. Interneurons and motor neurons regenerate after zebrafish SCI. Notably, dopamine and serotonin signals from regenerating tracts control motor neuron regeneration by promoting the proliferation of pMN ERGs (Barreiro-Iglesias et al., 2015; Reimer et al., 2013). We found *mstnb* mutants display increased neuronal differentiation at 1 wpi, with an overrepresentation in glycinergic interneurons among regenerating neurons. Glycinergic inhibition plays important roles in coordinating locomotor rhythms in different organisms (Hinckley et al., 2005; Jovanović et al., 1999; Sibilla and Ballerini, 2009). We propose that increased inhibitory neurotransmission disrupts E/I balance and may underlie the behavioral recovery defects observed in *mstnb* mutants. The mechanisms that underlie E/I balance disruption in *mstnb* mutants require further investigation into the time course of neuronal regeneration and the contribution of *mstnb^+^* ERGs to specific neuronal populations.

Our study highlights a niche of dorsal ependymal progenitors that express *mstnb* after SCI. Unlike tissues that undergo constant cell renewal such as skin or blood, the nervous system undergoes little turnover and does not harbor a constitutively active neurogenic niche. Instead, neural progenitors are quiescent and are only activated upon physiological or pathological stimulation. Lineage restricted ependymal progenitors emerge after zebrafish SCI. ERG niches include a ventro-lateral domain that gives rise to regenerating motor neurons, and a ventral domain that undergoes epithelial-to-mesenchymal transition and is required for glial bridging after SCI (Klatt Shaw et al., 2021; Reimer et al., 2008). Our findings support the emergence of a lineage restricted, neurogenic niche of dorsal ERGs during SC regeneration in zebrafish. Consistent with this model, the numbers of motor neurons and the extent of glial bridging across the lesion, which have been respectively associated with ventro-lateral and ventral ERGs, are unaffected in the absence of *mstnb*. Instead, *mstnb* mutants showed specific neurogenesis defects in dorsal SCs, and a preferential increase in dorsal glycinergic neurons. The molecular identity and cellular contributions of *mstnb^+^* ERGs to neurogenesis and SC repair warrant further investigation.

Niches of progenitor cells require intricate regulatory mechanisms to balance the rates of self- renewal, differentiation, and quiescence (Li and Clevers, 2010). At the cellular level, the organization of progenitor cells into localized niches maintains quiescence under homeostatic conditions, and triggers progenitor cell activation following niche disruption (Bagheri-Mohammadi, 2021). Molecularly, progenitor cell niches are hubs for Bmp, Wnt, and Notch signaling pathways, which control the rates of self-renewal, differentiation, and quiescence. Our study supports a model in which Mstn restricts neuronal differentiation and maintains neuronal progenitors in a proliferative, undifferentiated cell fate. Our findings are consistent with previously reported functions for Mstn in muscle, fat, and bone tissues (Dogra et al., 2017; Langley et al., 2002; Le and Yao, 2017; Lim et al., 2018; McCroskery et al., 2003; Wallner et al., 2017). Notably, previous findings have shown that Mstn is a regeneration limiting gene for zebrafish heart or fin regeneration, which are both dedifferentiation-based repair mechanisms (Dogra et al., 2017; Magga et al., 2019; Uribe et al., 2018). In contrast, in the context of SC regeneration, we find that Myostatin promotes regeneration by supporting regenerative FGF signaling, revealing a new role for Myostatin in this stem cell-based regeneration paradigm. Together these findings underline how tissue regeneration programs can coopt similar signaling pathways to achieve highly specific regenerative outcomes, and indicate a tissue-specific mechanism for Myostatin signaling.

We propose that Fgf is a mediator of Mstn functions during SC regeneration, and that Mstn limits neuronal differentiation by promoting Fgf-dependent self-renewal in *mstnb^+^* ERGs. Similar regulatory mechanisms have been shown in muscle tissues, where Mstn inhibits the muscle differentiation transcription factors MyoD and Myogenin. Fgf signaling promotes NSC proliferation and self-renewal (Hsu et al., 2009). In mammals, isolated ependymal cells have the capacity to form neurospheres and produce neurons, astrocytes and oligodendrocytes *in vitro* (Meletis et al., 2008). However, although mammalian SCI induces the proliferation of ependymal cells lining the central canal (Horner et al., 2000), mammalian ependymal cells are incapable of forming neurons *in vivo* (Barnabe-Heider et al., 2010; Muthusamy et al., 2018; Ren et al., 2017; Shah et al., 2018). We propose that comparative studies between zebrafish ERGs and mammalian ependymal cells could reveal new insights into their differential regenerative capacities and examine whether Mstn signaling is differentially regulated between zebrafish and mammals.

## ACKNOWLEDGMENTS

We thank V. Cavalli, A. Johnson, K. Poss, and L. Solnica-Krezel for discussion, D. Stainier for sharing *mstnb* mutants, T. Li and B. Zhang for Bioinformatics analysis, and the Washington University Zebrafish Shared Resource for animal care. This research was supported by grants from the NIH (R01 NS113915 to M.H.M.) and the McDonnell Center for Cellular Neuroscience (to M.H.M.).

## METHODS

### Zebrafish

Adult zebrafish of the Ekkwill, Tubingen, and AB strains were maintained at the Washington University Zebrafish Core Facility. All animal experiments were performed in compliance with institutional animal protocols. Male and female animals between 3 and 9 months of ∼2 cm in length were used. Experimental fish and control siblings of similar size and equal sex distribution were used for all experiments. SC transection surgeries and regeneration analyses were performed in a blinded manner, and 2 to 4 independent experiments were repeated using different clutches of animals. The following previously published zebrafish strains were used: *mstn^bns5^* (Dogra et al., 2017), Tg(*nes*:GFP) (Lam et al., 2009), Tg(*dbx1b*:GFP) (Satou et al., 2012), Tg(*slc6a5*:GFP) (McLean et al., 2007).

### SC transection and treatment

Zebrafish were anaesthetized using MS-222. Fine scissors were used to make a small incision that transects the SC 4 mm caudal to the brainstem region. Complete transection was visually confirmed at the time of surgery. Injured animals were also assessed at 2 or 3 dpi to confirm loss of swim capacity post-surgery. For sham injuries, animals were anaesthetized, and fine scissors were used to transect skin and muscle tissues without inducing SCI.

For pro-MSTN treatment, lyophilized human MSTN proform peptide (BioVision, 4623P-10) was reconstituted in ddH2O to a concentration 100 ng/μl. Zebrafish were anaesthetized using MS-222. 2 μl (200 ng) of reconstituted peptides were injected daily adjacent and lateral to the SC lesion site. 2 μl of ddH2O was injected for vehicle controls.

For FGF1 treatment, lyophilized human FGF1 protein (PeproTech, 100-17A) was reconstituted in heparin to a concentration 250 ng/μl. Sterile Gelfoam Absorbable Gelatin Sponge (Pfizer, 09- 0315-08) was cut into 2 mm^3^ pieces, soaked with 2 μl of recombinant FGF1, then cut into 10 smaller pieces (50 ng per piece). Vehicle gelfoam pieces were soaked with 2 μl of heparin solution. At 5 dpi, zebrafish were anaesthetized using MS-222 and longitudinal incision lateral and parallel to the SC was made with fine scissors. Injured SC tissues were exposed without causing secondary injuries and gelfoam sponges were places adjacent to the lesion site. The incision was closed and glued using Vetbond tissue adhesive material as previously described (Mokalled et al., 2016).

### Bulk RNA sequencing

Two mm SC sections, including the lesion site plus additional rostral and caudal tissue proximal to the lesion, were collected from *mstnb* mutants and wild-type siblings at 1 wpi. Uninjured *mstnb* mutants and wild-type SCs were also collected. Total RNA was prepared using NucleoSpin RNA Plus XS (Clontech, cat# 740990) and sent for bulk RNA sequencing. TruSeq libraries were prepared and sequenced on Illumina HiSeq 3000 using 50 bp single-end reading strategy. Quality QC and trimming of adapters and short sequences were performed using Fastx. Sequencing reads were mapped to the zebrafish genome (Zv11) using Bowtie2, then assembled and quantified using the Cufflinks and Cuffdiff algorithms. Genes with log2(fold enrichment) between -1 and 1 or adjusted p-value ≥ 0.01 were considered insignificant. RNA sequencing was performed at the Genome Technology Access Center at Washington University. Analysis was performed in the Bioinformatics Core at the Center for Regenerative Medicine at Washington University.

RNA-seq data (GEO accession number : GSE77025) was used to evaluate the expression of Tgf- β ligands after complete SC transection. Log2(fold change) is expressed for SCs at 1 wpi relative to the sham injured SCs (Mokalled et al., 2016).

### Histology

16 µm µm cross cryosections of paraformaldehyde-fixed SC tissue were used. Tissue sections were imaged using a Zeiss AxioVision compound microscope for *in situ* hybridization or a Zeiss LSM 800 confocal microscope for immunofluorescence. *In situ* hybridization for *mstnb* was performed as previously described (Mokalled et al., 2016).

For immunohistochemistry, tissue sections were rehydrated in PBT (0.1% Tween-20 in PBS), then treated with blocking agent (5% goat serum in PBT) for 1 hr at room temperature. For nuclear antigens, sections were treated with 0.2% TritonX-100 in PBT for 5 minutes and washed thoroughly in PBT prior to the blocking step. Sections were incubated overnight with the indicated primary antibodies, washed in PBT, and treated for 1 hr with secondary antibodies. Following washes, sections were incubated in 1 mg/mL of Hoechst and mounted in Fluoromount-G mounting media. Primary antibodies used in this study were: rabbit anti-Smad3(S423/425) (Abcam, ab52903, 1:50), rabbit anti-PCNA (GeneTex, GTX124496, 1:500), mouse anti-HuC/D (Invitrogen, A21271, 1:500), mouse anti-Gfap (ZIRC, Zrf1, 1:1000), mouse anti-acetylated α- tubulin (Sigma, T6793, 1:1000), chicken anti-GFP (Aves Labs, 1020, 1:1000), rabbit anti-Sox2 (GeneTex, 124477, 1:250). Secondary antibodies (Invitrogen, 1:200) used in this study were Alexa Fluor 488- or Alexa Fluor 594- conjugated goat anti-rabbit, anti-mouse, or anti-chicken antibodies.

For simultaneous labeling with rabbit anti-Sox2 (GeneTex, 124477, 1:250) and rabbit anti- pSmad3(S423/425) (abcam, ab52903, 1:50) (Fig. 1B), unconjugated Fab Fragment Goat Anti- Rabbit IgG(H+L) (Jackson ImmunoResearch, 111-007-003) and donkey anti-goat 568 (Thermo fisher: A-11057) antibodies were used for pSmad3 labeling. Sox2 was labeled using donkey anti- rabbit-488 (Jackson ImmunoResearch, 711-547-003).

EdU Staining was adapted from a a previously described protocol (Salic and Mitchison, 2008). Briefly, zebrafish were anaesthetized using MS-222 and subjected to intraperitoneal EdU injections. 12.5 mM EdU (Sigma 900584) diluted in PBS was used. A single injection (Fig. 3 and S2) or multiple, daily injections (Fig. 4 and S3) were performed and paraformaldehyde-fixed cryosections were used. Sections were rehydrated in PBT for 10 min then incubated with freshly prepared staining solution for 30 min (100 mM Tris (Sigma, T6066) pH 8.5; 1 mM CuSO4 (Sigma, C1297); 10 µM fluorescent azide; and 100 mM ascorbic acid (Sigma, A5960)).

### Cell counting

Cell counting was performed using a customized Fiji script (adapting ITCN: Image based Tool for counting nuclei- https://imagej.nih.gov/ij/plugins/itcn.html). Orthogonal projections of individual image stacks were generated using Zen software. A Customized Fiji script incorporated user-defined inputs to define channels (including Hoechst), to determine the center of the central canal, and to outline SC perimeters. SC tissues dorsal to the central canal center was considered “dorsal SC”. SC tissues ventral to the central canal center was considered “ventral SC”. To quantify nuclei, the following parameters were set in ITCN counter: width, 15; minimal distance, 7.5; threshold, 0.4. For each staining, thresholds were user-defined. Raw counts from Fiji were processed using a customized R Studio script.

### Swim endurance assays

Zebrafish were exercised in groups of 8-12 in a 5 L swim tunnel device (Loligo, cat# SW100605L, 120V/60Hz). After 10 minutes of acclimation inside the enclosed tunnel, water current velocity was increased every two minutes and fish swam against the current until they reached exhaustion. Exhausted animals were removed from the chamber without disturbing the remaining fish. Swim time and current velocity at exhaustion were recorded. Results were expressed as means ± SEM. An unpaired two-tailed Student’s t-test with Welch correction was performed using the Prism software to determine statistical significance of swim times between groups.

### Swim behavior assays

Zebrafish were divided into groups of 5 in a 5 L swim tunnel device (Loligo, cat# SW100605L, 120V/60Hz). Each group was allowed to swim for a total of 15 min under zero to low current velocities (5 min at 0 cm/s, 5 min at 10 cm/s, and 5 min at 15 cm/s). The entire swim behavior was recorded using high-speed camera (iDS, USB 3.0 color video camera) with following settings: aspect ratio, 1:4; pixel clock, 344; frame rate, 70 frames/s; exposure time: 0.29; aperture, 1.4 to 2; maximum frames; 63,000. Movies were converted to 20 frames/s and analyzed using a customized Fiji macro. For each frame, animals/objects > 1500 px^2^ were identified, and the XY coordinates were derived for each animal/object. Frame were independently, and animal/object tracking was completed using a customized R Studio script. The script aligned coordinates, calculated swim metrics considering three separate frame windows (Frames 0-6000 at 0 cm/s; frames 6001-12000 at 10 cm/s, and frames 12001-18001 at 20 cm/s).

### Glial bridging

GFAP immunohistochemistry was performed on serial transverse sections. The cross-sectional area of the glial bridge and the area of the intact SC rostral to the lesion were measured using ImageJ software. Bridging was calculated as a ratio of these measurements. Mann Whitney tests were performed using Prism software to determine statistical significance between groups.

### Axon tracing

Anterograde axon tracing was performed on adult fish at 4 wpi. Fish were anaesthetized using MS-222 and fine scissors were used to transect the cord 4 mm rostral to the lesion site. Biocytin-soaked Gelfoam Gelatin Sponge was applied at the new injury site (Gelfoam, Pfizer, cat# 09-0315-08; Biocytin, saturated solution, Sigma, cat# B4261). Fish were euthanized 6 hours post-treatment and Biocytin was histologically detected using Alexa Fluor 594-conjugated Streptavidin (Molecular Probes, cat# S-11227). Biocytin-labeled axons were quantified using the “threshold” and “particle analysis” tools in the Fiji software. Four sections per fish at 0.5 (proximal) and 2 (distal) mm caudal to the lesion core, and 2 sections 1 mm rostral to the lesion, were analyzed. Axon growth was normalized to the efficiency of Biocytin labeling rostral to the lesion for each fish. The axon growth index was then normalized to the control group for each experiment. Mann-Whitney tests were performed using Prism software to determine statistical significance between groups

### Quantitative real time PCR

Two mm SC sections, including the lesion site plus additional rostral and caudal tissue proximal to the lesion, were collected for qRT-PCR. Total RNA was prepared using NucleoSpin RNA Plus XS (Clontech, cat# 740990) and cDNA was synthesized using the Maxima First Strand cDNA Synthesis Kit (ThermoFisher, cat# K1672) according to manufacturer’s specifications. Quantitative PCR was completed using the Luna polymerase master mix (NEB, cat# M3003) using gene-specific primers (Table S1). Primers were designed to flank introns and were confirmed to not amplify project from genomic DNA. To determine primer efficiency, a standard curve was generated for each primer set using cDNA pooled from wild-type embryos at 1, 3, and 5 days post-fertilization. qRT-PCR was performed on a Bio-Rad CFX Connect Real-Time System. For each gene, log2(fold change) was calculated using the DCq method and normalized to *eif1a* as a loading control and to control gene expression for each experiment.

---

**Figure S1.**
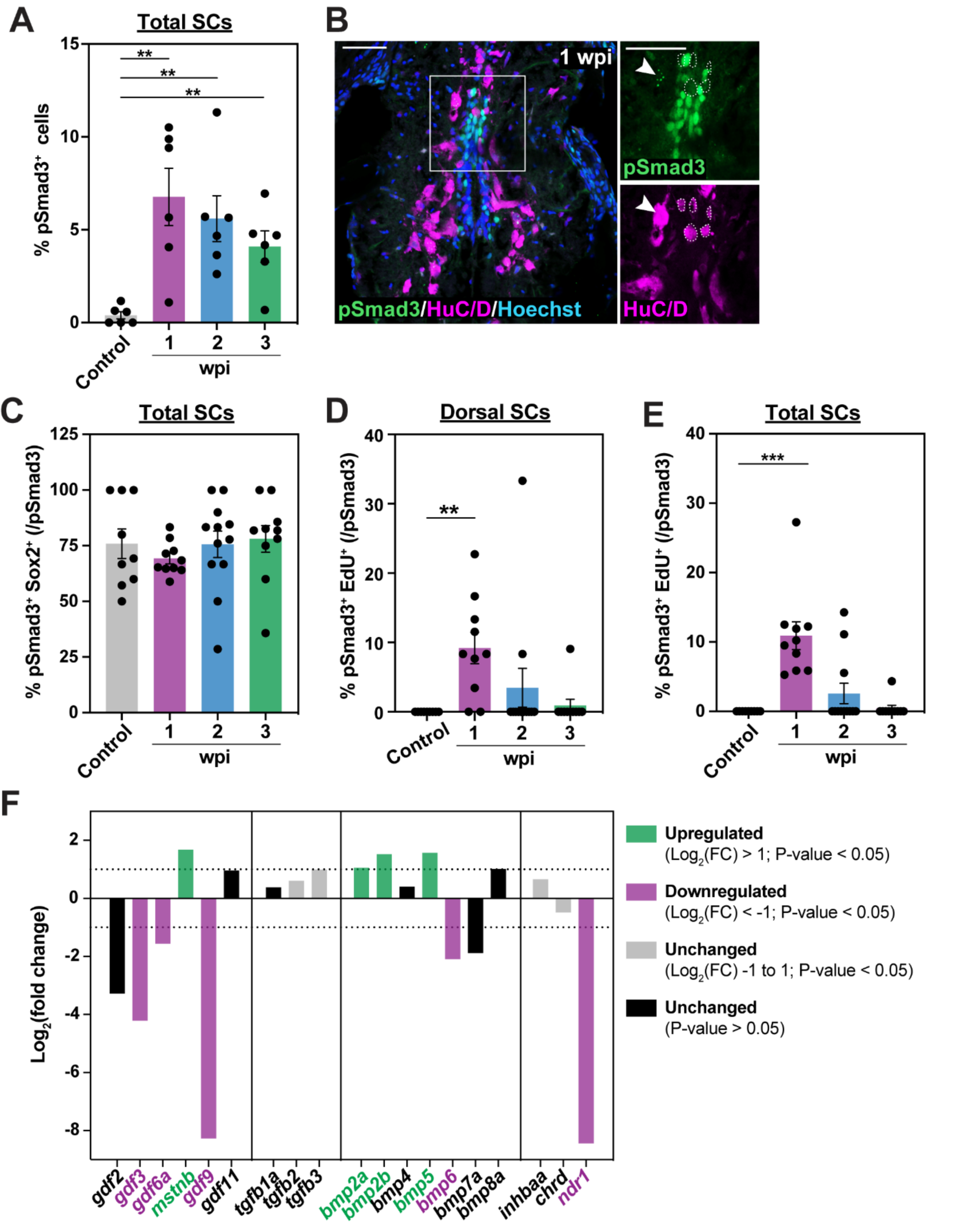
TGF-β signaling during SC regeneration. **(A)** pSmad3 quantification in total SC sections. Wild-type SCs at 1, 2 and 3 wpi and uninjured SCS were analyzed. Percent pSmad3^+^ cells was normalized to the number of nuclei in dorsal SCs. **(B)** pSmad3 and HuC/D immunostaining in wild-type SCs at 1 wpi. High-magnification views of dorsal SCs are shown. Dotted lines delineate pSmad3^-^ HuC/D^+^ neurons. Arrowheads point to dotted pSmad3 expression in a subset of HuC/D^+^ neurons. **(C)** pSmad3 quantification in total ERGs. Wild-type SCs at 1, 2 and 3 wpi and uninjured SCS were analyzed. Percent pSmad3^+^ Sox2^+^ cells was normalized to the number of pSmad3^+^ cells. **(D, E)** pSmad3 and EdU quantification in dorsal (D) and total (E) SCs. EdU was administered for 24 hrs prior to SC collection. Wild-type SCs at 1, 2 and 3 wpi and uninjured SCS were analyzed. Percent pSmad3^+^ EdU^+^ cells was normalized to the number of pSmad3^+^ cells. **(F)** Expression of Tgf-β ligands in wild-type SCs at 2 wpi by bulk RNA sequencing. For each gene, log2 (fold change) was normalized to gene expression levels in sham injured SCs. Upregulated (green) and downregulated (magenta) genes are shown. Unchanged genes are shown in grey and black. For all quantifications, cross SC sections 450 μm rostral to the lesion site were quantified. ***P<0.001; **P<0.01. Scale bars, 50 µm.

**Figure S2.**
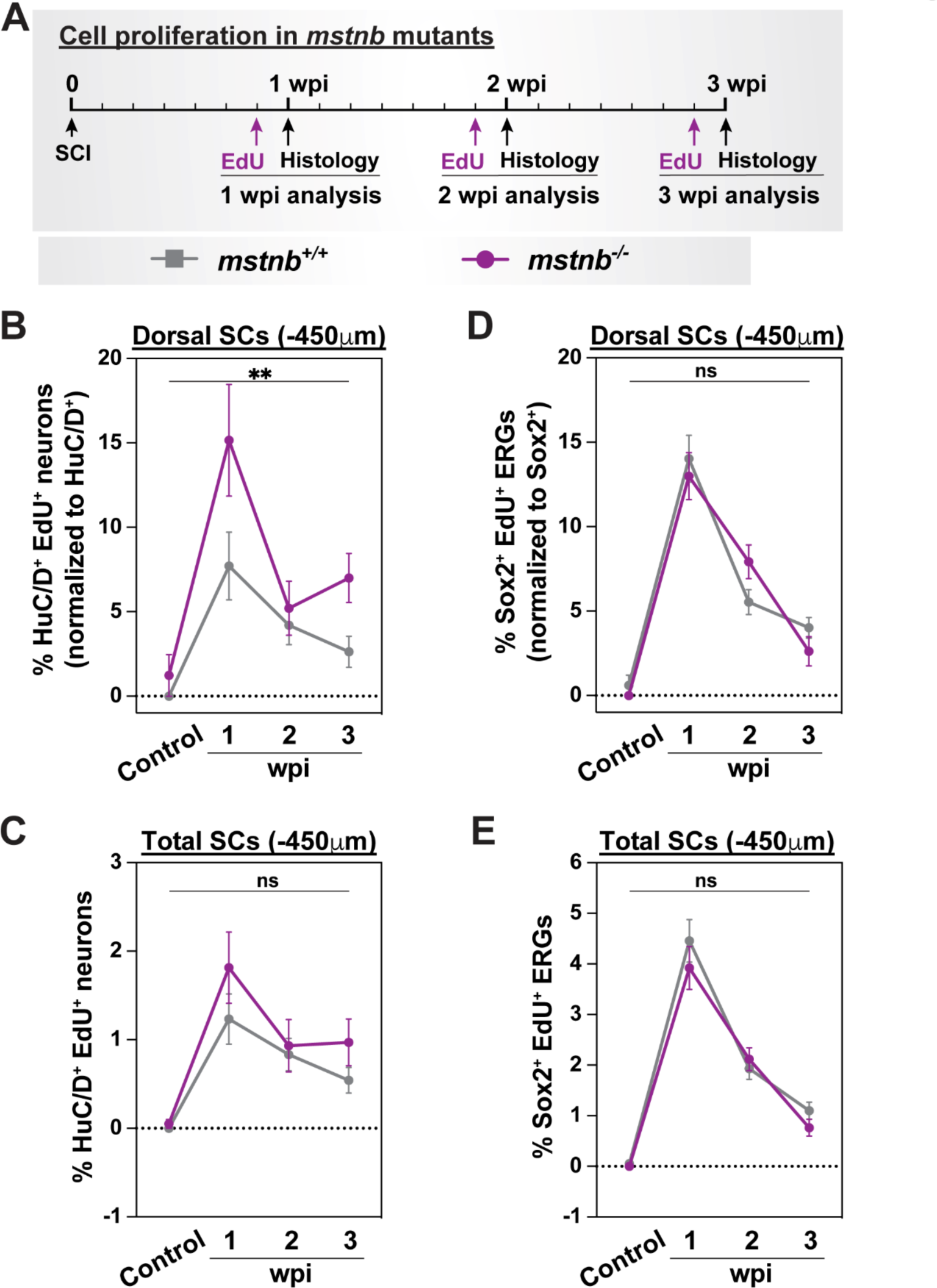
Cell proliferation in *mstnb* mutant zebrafish. **(A)** Experimental timeline to assess the rates of cell proliferation. *mstnb^-/-^* and wild-type siblings were subjected to SC transections. A single EdU injection was performed at either 6, 13, or 20 days post-injury. SC tissues were harvested for analysis at 1, 2, 3 wpi and at 24 hours after EdU injection. Animal numbers are indicated for each genotypes and two independent replicates are shown. **(B)** Regenerated HuC/D^+^ EdU^+^ neurons were quantified in dorsal SC sections at 1, 2, 3 wpi and uninjured controls. Percent HuC/D^+^ EdU^+^ neurons was normalized to the total number of HuC/D^+^ neurons for each section. **(C)** Regenerated HuC/D^+^ EdU^+^ neurons were quantified in total SC sections at 1, 2, 3 wpi and uninjured controls. Percent HuC/D^+^ EdU^+^ neurons was normalized to the total number of nuclei for each section. **(D)** Sox2^+^ EdU^+^ ERGs were quantified in dorsal SC sections at 1, 2, 3 wpi and uninjured controls. Percent Sox2^+^ EdU^+^ ERGs was normalized to the total number of Sox2^+^ ERGs for each section. **(E)** Sox2^+^ EdU^+^ ERGs were quantified in total SC sections at 1, 2, 3 wpi and uninjured controls. Percent Sox2^+^ EdU^+^ ERGs was normalized to the total number of nuclei for each section. For all quantifications, cross SC sections 450 μm rostral to the lesion site were quantified.**P<0.01; ns, not significant.

**Figure S3.**
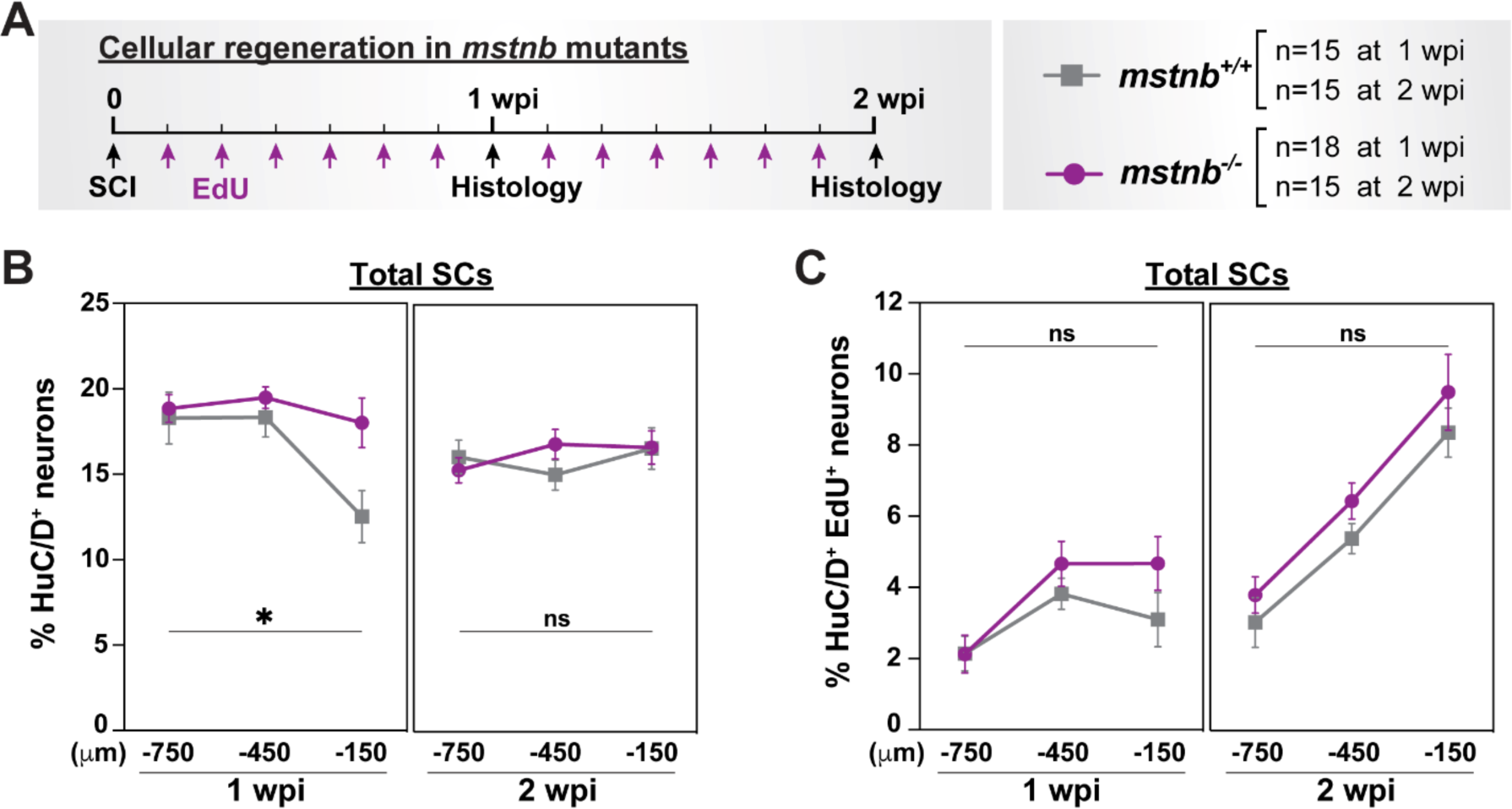
Regenerative neurogenesis in *mstnb* mutant zebrafish. **(A)** Experimental timeline to assess the rates of neurogenesis and ERG self-renewal. *mstnb^-/-^* and wild-type siblings were subjected to SC transections and daily EdU injections. SC tissues were harvested for analysis at 1 or 2 wpi. Animal numbers are indicated for each genotypes and two independent replicates are shown. **(C)** HuC/D^+^ neurons were quantified in total SC sections at 1 and 2 wpi. Percent HuC/D^+^ neurons was normalized to the total number of nuclei for each section. **(D)** Regenerated HuC/D^+^ EdU^+^ neurons were quantified in total SC sections at 1 and 2 wpi. Percent HuC/D^+^ EdU^+^ neurons was normalized to the total number of nuclei for each section. For all quantifications, cross SC sections at 150, 450, and 750 μm rostral to the lesion site were quantified. **P<0.01; ns, not significant.

**Figure S4.**
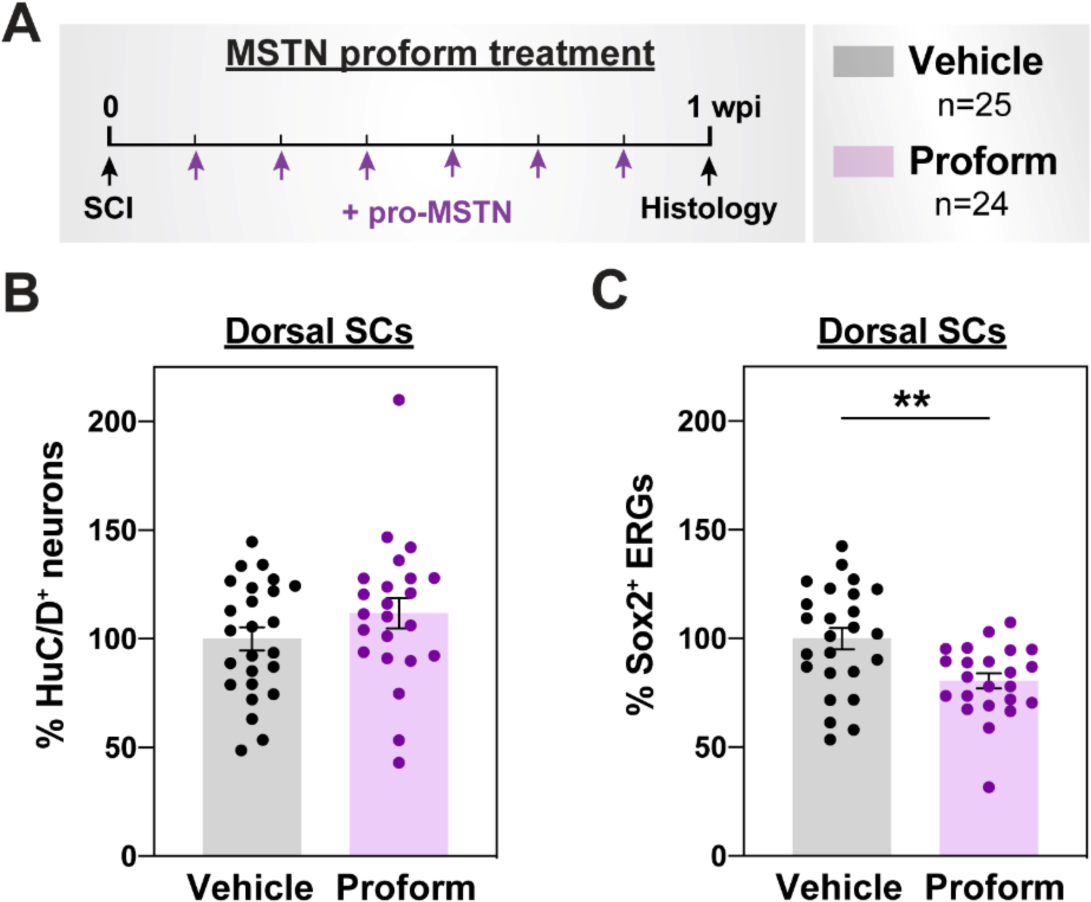
Pharmacological Mstn inhibition during SC regeneration. **(A)** For local Mstnb inhibition, wild-type SCs were subjected to SC transections and daily injections of human recombinant MSTN Proform (pro-MSTN) peptide adjacent to the lesion site. SC tissues were harvested for analysis at 1 wpi. Animal numbers are indicated for each genotypes and two independent replicates are shown. **(B)** HuC/D^+^ neurons were quantified in pro-MSTN- and vehicle-treated SCs. Dorsal SC sections at 1 wpi were analyzed. Percent HuC/D^+^ neurons was normalized to the total number of nuclei for each section. **(C)** Sox2^+^ ERGs were quantified in pro- MSTN- and vehicle-treated SCs. Dorsal SC sections at 1 wpi were analyzed. Percent Sox2^+^ ERGs was normalized to the total number of nuclei for each section. For all quantifications, cross SC sections 450 μm rostral to the lesion site were quantified. **P<0.01.

**Figure S5.**
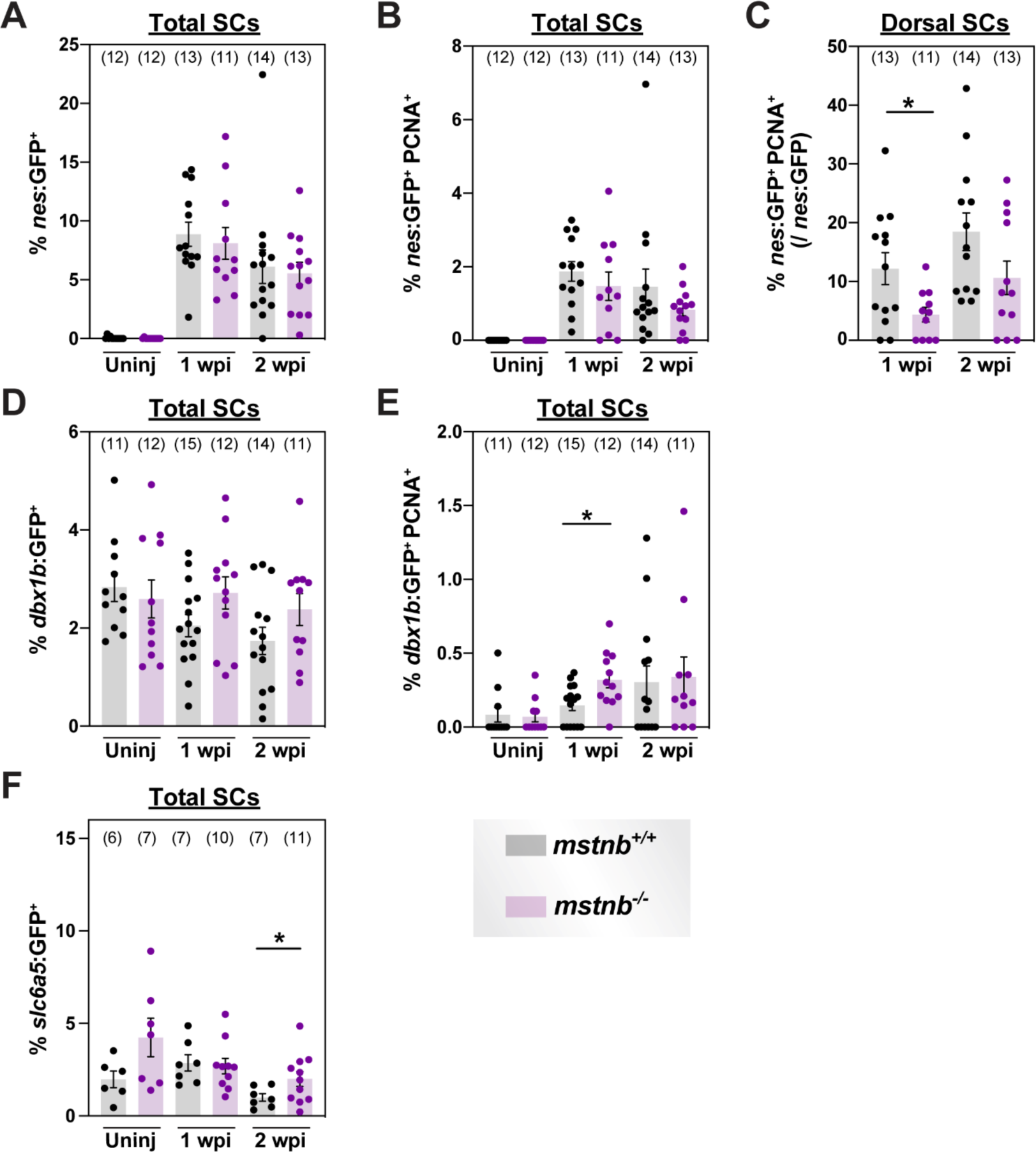
Assessment of neuronal progenitors and neurons in *mstnb* mutant zebrafish. **(A)** *nes^+^* NSCs were quantified in total SC sections. Percent *nes^+^* NSCs was normalized to the total number of nuclei for each section. **(B)** *nes^+^* PCNA^+^ NSCs were quantified in total SC sections. Percent *nes^+^* PCNA^+^ NSCs was normalized to the total number of nuclei for each section. **(C)** *nes^+^* PCNA^+^ NSCs were quantified in dorsal SC sections. Percent *nes^+^* PCNA^+^ NSCs was normalized to the total number of *nes^+^* NSCs for each section. **(D)** *dbx1b^+^* iNPs were quantified in total SC sections. Percent *dbx1b^+^* iNPs was normalized to the total number of nuclei for each section. **(E)** *dbx1b^+^* PCNA^+^ iNPs were quantified in total SC sections. Percent *dbx1b^+^* PCNA^+^ NSCs was normalized to the total number of nuclei for each section. **(F)** *slc6a5*:GFP*^+^* glycinergic neurons were quantified in total SC sections. Percent *slc6a5*:GFP*^+^* neurons was normalized to the total number of nuclei for each section. For all quantifications, cross SC sections 450 μm rostral to the lesion site were quantified. *P<0.05.

**Table S1.**
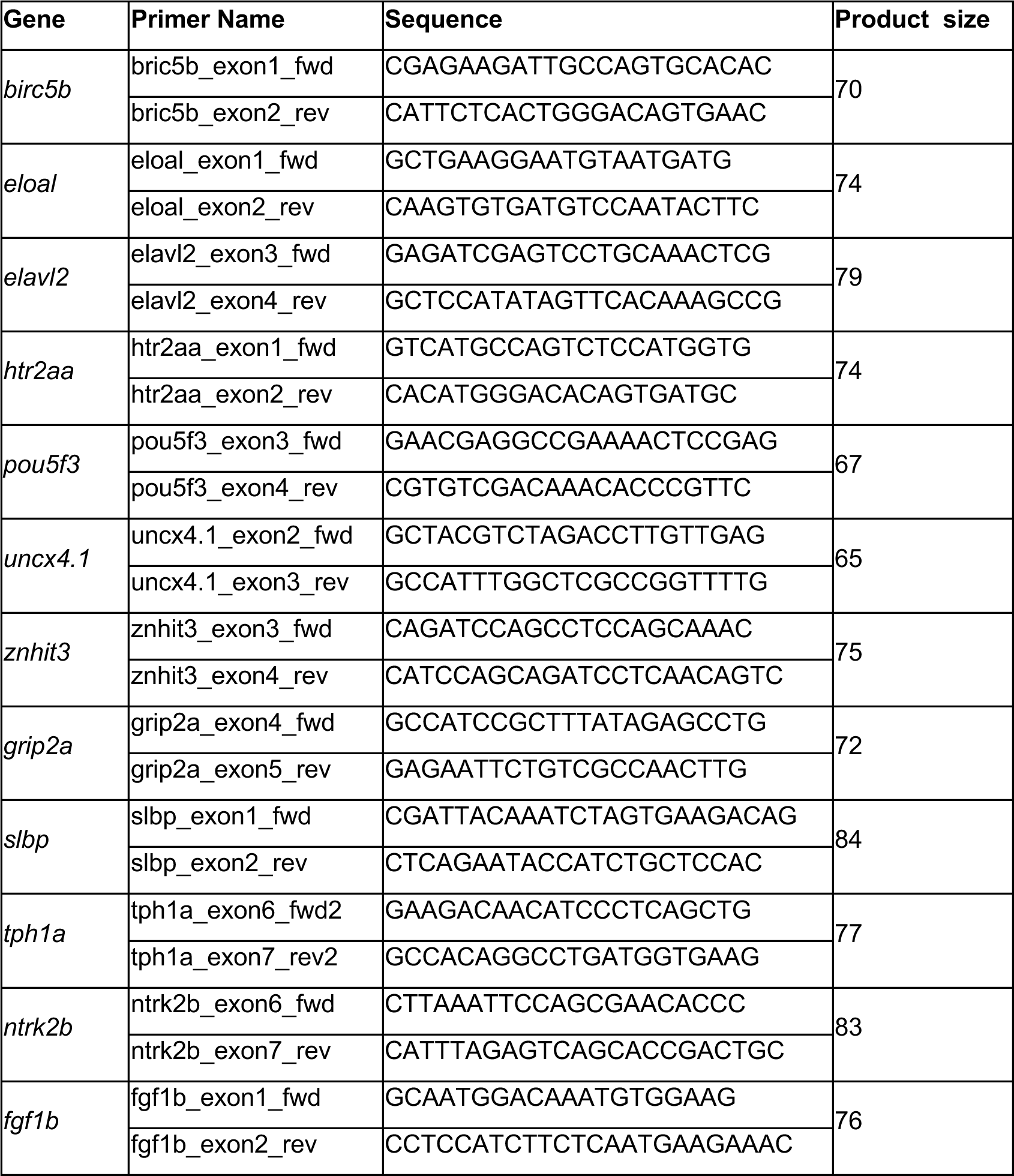
Primer sequences for qRT-PCR. Gene names, primer names, sequences, and product sizes are indicated.

